# RAFTS^3^G – An efficient and versatile clustering software to analyses in large protein datasets

**DOI:** 10.1101/407437

**Authors:** Bruno Thiago de Lima Nichio, Aryel Marlus Repula de Oliveira, Camilla Reginatto de Pierri, Leticia Graziela Costa Santos, Ricardo Assunção Vialle, Jeroniza Nunes Marchaukoski, Fabio de Oliveira Pedrosa, Roberto Tadeu Raittz

## Abstract

The need to develop computational tools and techniques that can predict efficiently consistent groups of family proteins in large volume of biological information is still a great perspective in Bioinformatic studies. Besides that, it is difficult to increase speed demanding low computational processing to minimize the information complexity. Tools already consolidated as the CD-HIT and UCLUST generates very compact data that makes the Data Mining difficult and have low efficiency when used for detect homology among proteins requiring manual intervention, therefore it is necessary a tool that is also efficient in low similarity. Here we present a new approach for the Data Mining and analysis of homology in large dataset of protein sequences, the RAFTS^3^G. We used the UniProtKB/Swiss-Prot database with the most popular clustering tools and RAFTS^3^G proved to be more than 10 times faster than CD-HIT and its strategy increases the performance in low similarity to detect protein families.

## 1 Introduction

Since the emergence of large-scale genomic sequencing, in 2002, the analyses of genomes and proteomes began to be used and gained strength, mainly in recent years, however, it was noted that there was an exponential increase of more and more sequences to be deposited resulting in the necessity of creating large databases to store such information what we call Big Data [1]. Such growth was responsible for bringing a major bottleneck to the field of analysis of sequences: the need to develop computational tools and techniques that can handle large volume of biological information that minimize the redundancy [2]. Another problem is the application of a high level of programming knowledge on the part of researchers to analyze large volumes of data, which hinders the fluidity of the researches [3]. Nowadays two tools are very relevant in the clustering sequences to minimize redundancy in large dataset: CD-HIT and UCLUST. CD-HIT is one of the most popular tools and is the state-of-art method [4] [5]. UCLUST is a tool used by thousands of users around the world as high-performance clustering being faster than the CD-HIT algorithm [6]. However, CD-HIT does not support values lower than 40% of similarity and in lower identities, UCLUST is questionable because it degrades the quality of alignment [4]. Therefore, both CD-HIT and UCLUST are not reliable tools for parsing with homologies in conserved structures in large datasets with values less than 30% of similarity [7]. The study of homology is a great field of computational process uses, statistical detection of logical or in evolutionary analyses but building relationships between genomes or proteomes requires a lot of effort and time, mainly in large datasets [8]. The most efficient techniques in homology prediction use as gold standard the Basic Local Alignment Search Tool (BLAST) ‘all-against-all’ [3] or, in another cases, Markov Clustering (MCL) method adaptations [9], however it is dependent on alignment metrics requiring a lot of processing and time to generate results.

Alignment-free methods are strong alternatives to alignment-dependent techniques and are also efficient in minimizing the redundancy of biological data. These methods are computationally fast and use less memory compared to alignment-based methods and benchmark data sets are needed to explore the full potential of alignment-free methods [10].

These methods have been applied to problems ranging from whole-genome to the classification of protein families and the strength of these methods makes them particularly useful for Next-Generation Sequencing (NGS) data processing and analysis [11]. In order to explore the potential of the alignment free method associated with a strategy similar to BLASTP ‘all-against-all’, we developed the RAFTS^3^G, a new alternative proposal to clustering analyses no loss of biological data important to the homology study, easier for the researcher, requiring less processing, consequently being faster, even dealing with large volumes of biological data.

## Methods

### RAFTS^3^G strategy

As an easy way to the user, in order to minimize time and maintaining consistency in data analysis with proteins, we developed *Rapid Alignment Free Tool for Sequences Similarity Search to Groups* (RAFTS^3^G) tool. RAFTS^3^G was written in MATLAB v2017a using its built-in functions, the Bioinformatics Toolbox and an in-house library - including RAFTS3 algorithm (**Figure 3, supplementary**). As the name implies, the tool minimizes the use of alignment calculations because it is an alignment-free tool that searches for similarities between protein sequences from FASTA database and uses a filter *Hash* function for the selection of candidates is based on *k-mers* shared and a measure of comparison using a co-occurrence matrix of amino acid residues (BCOM) (as default, 18 *k-mers* with lengths of 10 amino acid residues are selected per sequence) [12]. RAFTS^3^G receive as input sequences file in multi FASTA format to be grouped. The script creates a base in which the query is performed by the RAFTS^3^ algorithm. Each protein in the input file is evaluated, initially, as a group, and the result of the query to the base assesses the similarity between the proteins by the self-score alignment value.

The RAFTS^3^G functioning takes place in two main stages: the first performs search of proteins in formatted database using RAFTS^3^ pipeline. The second stage RAFTS^3^G mounts the clusters through based on “all-versus-all” method under the similarity threshold. To the tests we adopted the threshold 0.5 (or 50%) of self-score analogous to 50% similarity implementing PAM and BLOSUM matrices. The clustering is analogous a BLASTP “all-against-all” method (**Figure 1)**. This strategy is the differential to clustering and consolidate homolog groups even low similarity among sequences, very important in homology analyses. The traditional BLAST “all-against-all” heuristics demands high computational power but it is very effective to detect homolog groups. RAFTS^3^G has as main feature complementary, run even with low computational requirements and in high or low similarities in large dataset of sequences (**Table 1, supplementary**).

**Figure 1.**
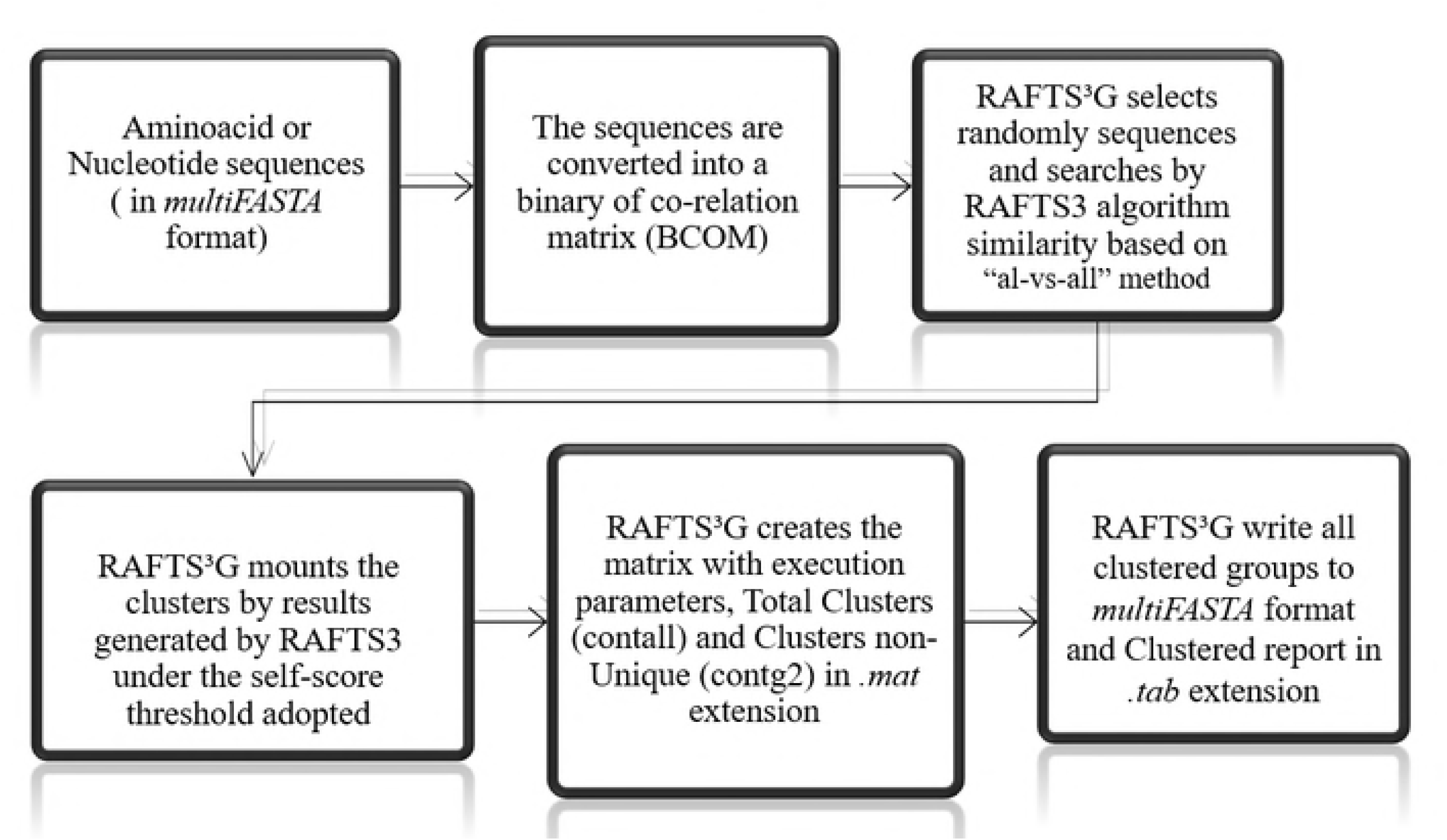
RAFTS^3^G workflow: A multiFASTA input is required and the sequences are converted to a binary matrix of amino acids. The RAFTS3 algorithm randomly selects the sequences and compares “all-against-all” and assembles the clusters with the similarity threshold. An matrix is generated with the execution data, cluster information and sequences grouped in .*mat* extension and all clusters generated are written in FASTA file and a log is generated in the .*tab* format.

## Results and discussion

### RAFTS^3^G performance in large dataset of proteins with low computational requirements

We selected seven different self-scores with of threshold 30, 40, 50, 60, 70, 80, 90 against the UniProt/Swiss-Prot in Machine 2 conditions (Table 1, supplementary). We observed that the cluster generation time was directly proportional to the increase in self-score. Having minimum clustering of about 5.8 hours in 30 self-score threshold and the time spent on self-score clustering with 90 total time of 13.9 hours. It was noted also that the average increase of unique clusters increased by 0.23 and 0.13 in representative clusters and clusters of approximately 0.20 on Total clusters. In relation to the time the tool kept an average of 10.75 hours, as shown in **Figure 3**.

**Table 1.** Correlation ratio for 50 representative *clusters* analysis randomly selected generated using CD-HIT, UCLUST and RAFTS^3^G with 90%, 95% and 99% similarity with UniProt/Swiss-Prot database. The match represents the RAFTS^3^G identified proteins compared to CD-HIT and UCLUST.

|  | 90% |  | 95% |  | 99% |  |
| --- | --- | --- | --- | --- | --- | --- |
|  | UCLUST | CD-HIT | UCLUST | CD-HIT | UCLUST | CD-HIT |
| **RAFTS<sup>3</sup>G** | 0.90 | 0.91 | 0.88 | 0.92 | 0.98 | 0.98 |

### Software benchmark: CD-HIT, UCLUST and RAFTS^3^G

CD-HIT [13] and UCLUST [6] are the highlights in high performance in large numbers of data of proteins and then were used in comparative analysis with RAFTS^3^G against the UniProt/Swiss-Prot Database (updated until june, 2018) [14]. According to results using Swiss-Prot database in different self-scores, we can report that RAFTS^3^G can be used even in computers with little processing and computational resources even with large amounts of information (**Table 1, supplementary**). For the time analysis, under the same conditions of Hardware, the CD-HIT algorithm was that it took longer, more than 1000 times and 11 times of the UCLUST and RAFTS^3^G, respectively (**Figure 4** and **Table 2, supplementary**). It is also worth pointing out that both the CD-HIT and UCLUST tools require a preprocessing step in which the data to be rotated by the algorithms must be organized in order of sequence size, because both algorithms select the largest to minor sequences to choose the representative sequence to the group and align the others from them, not being a random process. The output formats of the three tools are different. While CD-HIT (**Figure 1, supplementary**) and UCLUST (**Figure 2, supplementary**) bring the user information about clusters generated, percentage between the grouped sequences and headers of the grouped sequences, RAFTS^3^G brings the output in FASTA format with the note of the generated clusters added in headers. It is ideal for users who want to rescue information from the integral sequences without having to re-analyze or re-align each cluster in order to study them.

**Table 2.**
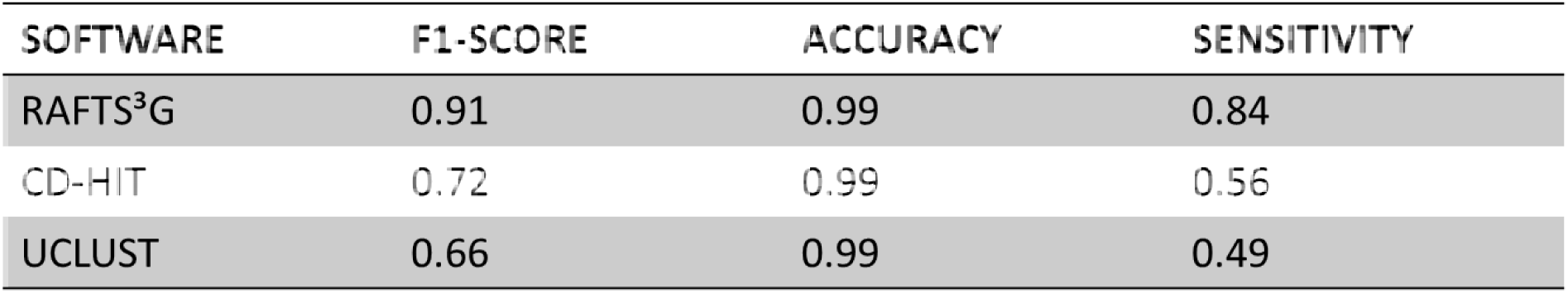
Software result with record the best F1-Score in relation to the cured families and presented the most relevant external metrics against Astral/SCOPe datasets in genetic domain sequence subsets with less than 95% identity to each other (release 2.07) using 50% up to 90% of similarity threshold.

**Figure 2.**
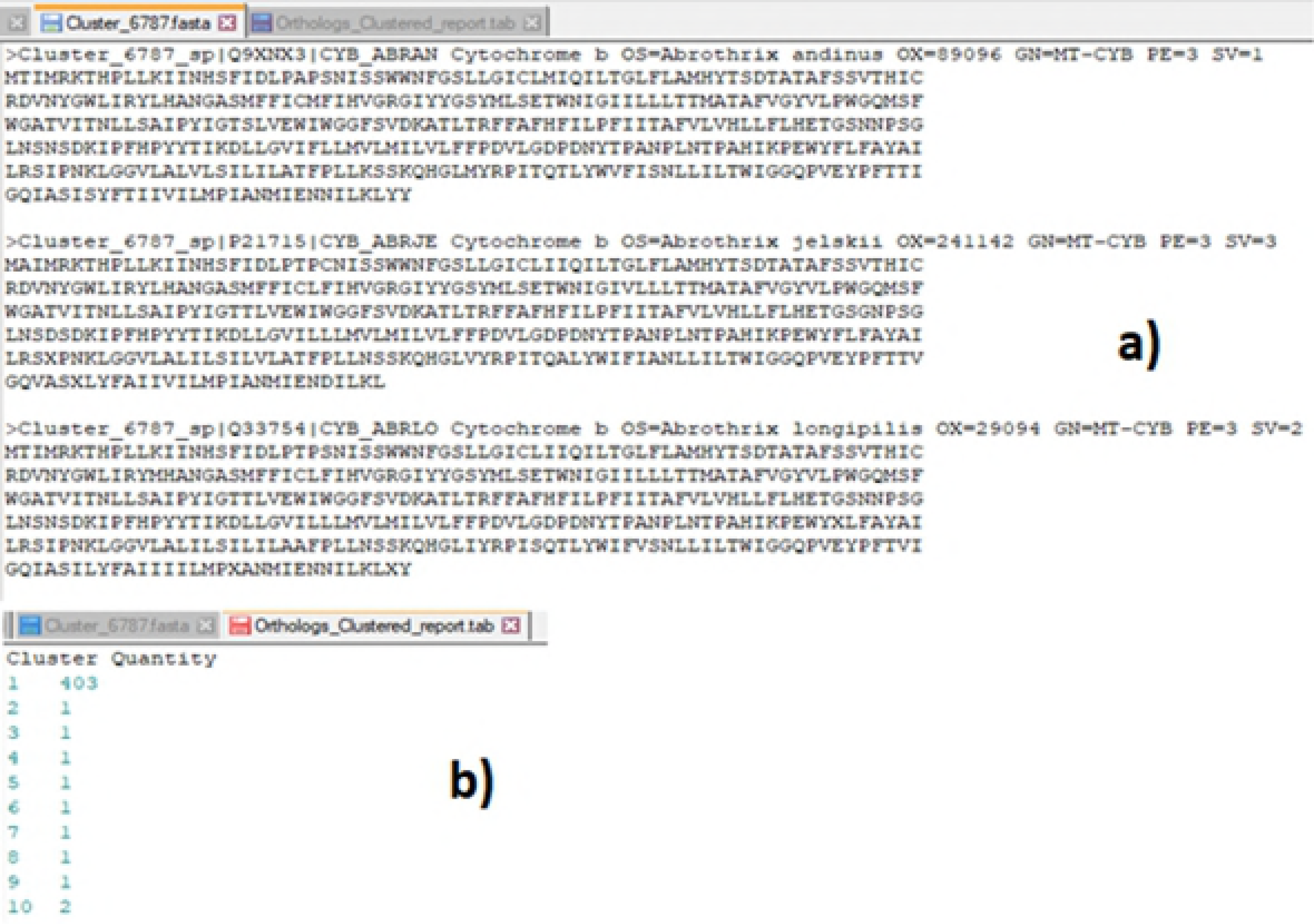
RAFTS^3^G outputs. An example of RAFTS^3^G clustered sequences in FASTA format **a)** and the clustered report in tab extension **b)**.

**Figure 3.**
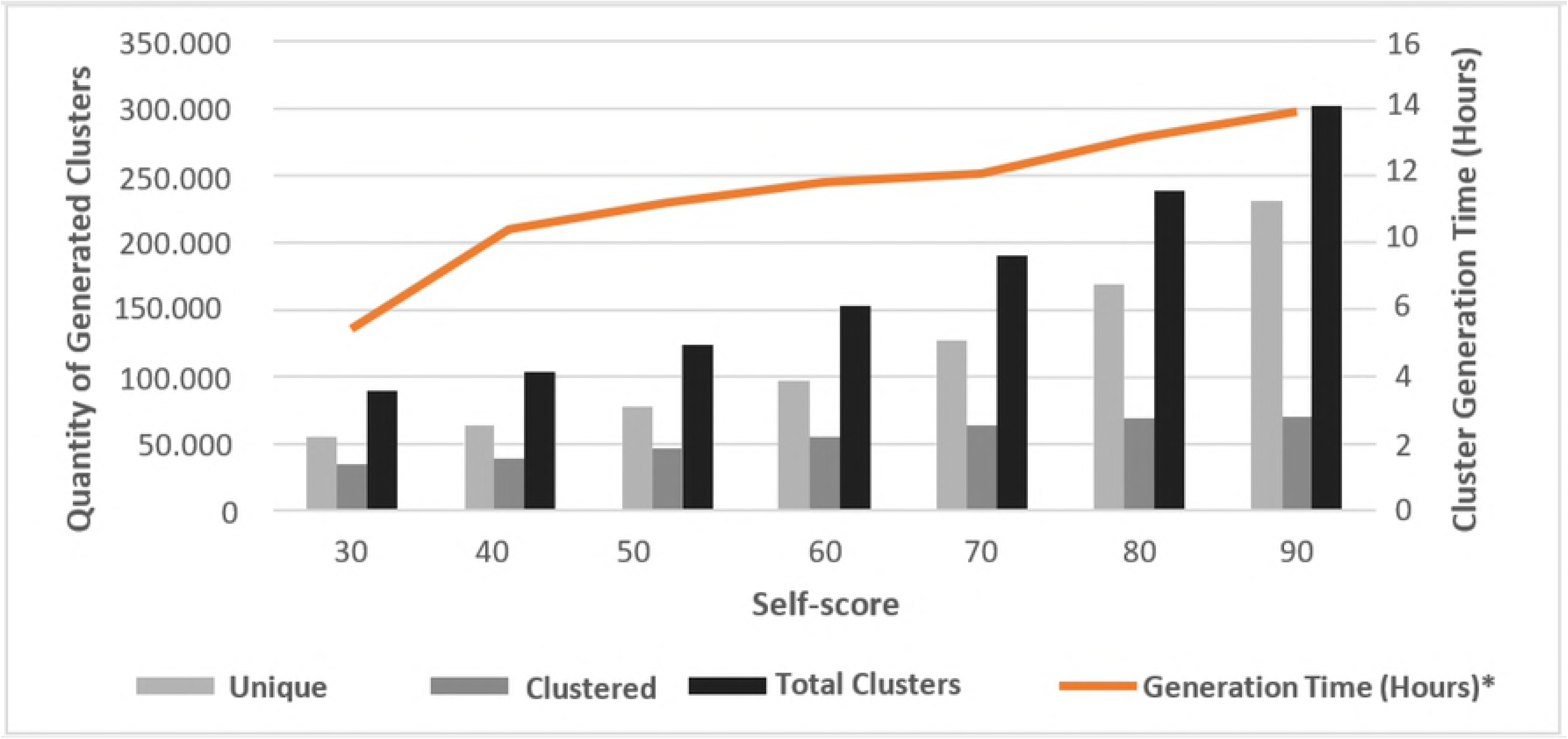
Unique sequences, Clustered sequences and Total Clusters (Unique + Clustered) Generated by the RAFTS^3^G tool in different self-scores in Machine 1 conditions. Relationship between time in hours of analysis (the vertical line on the right), number of Clusters generated (vertical line) in 7 different self-scores (horizontal line). * The value includes, besides the clustering algorithm, the formatting of the binary matrix Format Database.

### Clustering comparisons: CD-HIT, UCLUST and RAFTS^3^G

In high similarity – we adopted 90% 95% and 99% threshold - the study revealed that RAFTS^3^G, compared to CD-HIT and UCLUST in same conditions and set of data, obtained a little more sequences by cluster (**Table 3, supplementary**). This is possible because the algorithm selects randomly all proteins in the dataset to representative, differently in CD-HIT and UCLUST that selects representative sequences -the biggest protein sequences- and the clusters are generated from them. Despite this, the clusters generated by RAFTS^3^G approached from the other two tools in high similarity in up to 98% of the observed data - as shown in the **Table 1**.

### RAFTS^3^G homolog protein detector

With the tests in low similarities, RAFTS^3^G is the better choice for homology analysis. The RAFTS^3^G tool presented more than 0.90 sensitive in the grouping sequences belonging to the same family in 50% similarities against 0.79 of CD-HIT and 0.58 UCLUST algorithms (**Figure 5, supplementary**) and with 40% similarity RAFTS^3^G presented approximately 0.83 sensitive against 0.50 of CD-HIT and 0.49 UCLUST (**Figure 6, supplementary**).

The results are strengthened when RAFTS^3^G was performed using Astral/SCOPe datasets in genetic domain sequence subsets with less than 95% identity to each other. The three tools present the same accuracy 0.99. RAFTS^3^G highlights when observed sensitivity and F1-Score parameters, 0.84 and 0.91 respectively, -as shown in **Table 2.**

These results corroborate with the proposal of the CD-HIT and UCLUST, which is to optimize the accuracy to the detriment of the sensitivity to gain speed while maintaining the consistency of the clusters even if they become incomplete - lower sensitivity -, and therefore, the metrics reflect their goal. The RAFTS^3^G is more permissive for entering sequences in the clusters compared to the CD-HIT and USEARCH and therefore increased the sensitivity in this case. The three algorithms are different proposals, but that RAFTS^3^G is better for generating clusters for analysis and data mining and in homolog proteins detection because it loses much less related sequences compared to the CD-HIT and UCLUST, by the definition of the strategy.

### Conclusion and perspectives

The goal of this study is to bring a fast and efficient tool for the analysis of clusters involving large volume of data with low computational requisites introducing a new clustering approach using the alignment-free algorithm. RAFTS^3^G is comparable in more than 94% of the data generated by CD-HIT and UCLUST in high similarity, demonstrating its potential in reducing the redundancy of biological data. RAFTS^3^G is based on the BLASTP ‘all-against-all’ strategy and this contributes to the improvement of detection of homolog groups in lower similarity. RAFTS^3^G is about 11 times faster than CD-HIT algorithm and in lower similarities RAFTS^3^G is 0.22 and 0.33 more sensitive than CD-HIT and UCLUST respectively in detection of protein families. The data generated is easier to be rescued by the end user because it is in FASTA format with an extra log is generated to support the analyses. RAFTS^3^G also shows a great tool in the search for homologous proteins reaching 0.99 of accuracy and 0.84 sensitivity in homolog validated experimentally dataset. In addition, we performed RAFTS^3^G (50%) in NCBI NR database and we obtained 12,594,179 clusters composed of 4,127,885 non-Unique clusters and 8,466,294 Unique clusters of proteins. Therefore, with these studies we bring RAFTS^3^G how a new and versatile alternative to minimize redundancy in high similarities and detect homolog groups in large datasets, which is very important to comparative genomics, in NGS and sequence analyses under bioinformatics studies.

### Data and RAFTS^3^G availability

RAFTS^3^G is freely available at: https://sourceforge.net/projects/rafts-g/ Designed in Matlab v2017a and MCR runtime (v7.17) is required to runs. More information detailed in supplementary material.

## Acknowledgements

CAPES (Coordination for the Improvement of Higher Level -or Education-Personnel) & Araucária Fundations to support this work.

